# Near-critical slow dynamics enable flexible temporal computations and generalization

**DOI:** 10.64898/2026.06.29.735180

**Authors:** Gayathri Ramesan, Akhilesh Nandan, Daniel Koch, Aneta Koseska

## Abstract

Although neural activity often evolves along low-dimensional manifolds, such descriptions do not explain the dynamical mechanisms that generate, constrain, and stabilize computation. Identifying these mechanisms is essential for predicting responses to perturbations, understanding generalization to untrained signals, and explaining how similar computations arise from distinct circuit implementations. Here we use recurrent neural networks trained on an interval timing task as a model system to uncover the dynamical mechanisms of neural computation. We show that, despite converging to highly diverse attractor architectures, trained networks share a conserved transient dynamics. During learning, networks self-organize near dynamical bifurcations, forming structured ghost sets of slow points characterized by graded spectra of near-zero eigenvalues. These slow sets form a dynamical scaffold that constrains trajectory evolution. Inputs transiently reconfigure the vector field and reposition activity within this scaffold, while the underlying slow set governs subsequent dynamics. As a result, temporal computation is implemented through structured transient evolution rather than convergence to fixed points or persistent activity states. The extent of the slow sets predicts generalization to unseen temporal intervals, and networks lacking such organization fail to extrapolate reliably. To test sufficiency, we construct a minimal dynamical system endowed with analogous slow set geometry that reproduces interval timing without learning, providing a benchmark for identifying the essential dynamical ingredients of temporal computation. Together, these results identify structured slow transients as a candidate dynamical mechanism for temporal computation, provide a mechanistic interpretation of slow low-dimensional manifolds as emergent consequences of underlying state-space structure, and suggest that computational capacity in near-critical systems arises from the organization of transient flow rather than attractor states alone.

## Introduction

How neural populations perform computations through their dynamics remains a fundamental challenge in neuroscience. Across perception, memory, decision-making and action, behavior emerges from the transformation of information through precisely organized patterns of population activity evolving over time. Understanding neural computation therefore requires identifying the dynamical principles that generate, constrain and control these activity trajectories. Recent work has shown that neural population activity often evolves along low-dimensional trajectories Khona and Fiete (2022); Langdon et al. (2023); Perich and Gallego (2025). In interval timing tasks, in particular, neural dynamics occupy low-dimensional manifolds whose geometry and traversal speed correlate with behavioral output (Jazayeri and Afraz, 2017; Wang et al., 2018; Remington et al., 2018; Egger et al., 2019; Saxena and Cunningham, 2019; Bi and Zhou, 2020; Jazayeri and Ostojic, 2021; Beiran et al., 2023). Similar low-dimensional structures emerge in recurrent neural networks trained on analogous tasks, suggesting that they reflect general principles of neuronal computation (Wang et al., 2018; Remington et al., 2018; Beiran et al., 2023). These observations have motivated a geometric view in which computation is associated with low-dimensional manifolds, with task outputs encoded by positions or velocities along the manifold.

However, this framework leaves a fundamental mechanistic question unresolved: what dynamical structures generate and regulate these trajectories? Identifying a manifold describes where neural activity lies, but not the mechanisms that constrain the temporal evolution and determine its timescales. This distinction is important because the same low-dimensional trajectory can, in principle, arise from fundamentally different underlying dynamics, leading to different predictions about robustness, perturbation responses and generalization. Understanding computation therefore requires moving beyond geometric descriptions of neural activity to identify the dynamical mechanisms that generate it. This challenge is particularly acute for temporal computations, which are inherently transient Buonomano and Laje (2010); Zhou et al. (2022). Rather than converging to stationary attractors, neural populations must transform inputs through precisely organized patterns of activity that unfold over time. The central question is therefore not simply which low-dimensional manifolds exist, but which state-space structures organize transient evolution and thereby implement the computation itself.

Here we address this question by analyzing recurrent neural networks trained on an interval reproduction task and tracking the evolution of their dynamics during learning. We find that networks achieving similar behavioral performance converge to highly diverse attractor architectures, yet consistently develop a common transient dynamical organization. Across solutions, learning positions network dynamics near bifurcations, generating structured ghost-like sets of slow points (Strogatz and Westervelt, 1989) characterized by graded spectra of near-zero eigenvalues (Koch et al., 2024). These slow sets constrain the flow of the activity trajectories and organize their temporal evolution, providing a mechanism for reliable transient computation. Thus, while solutions are degenerate at the level of attractors, they are conserved at the level of transient dynamical organization. We further show that the computation emerges from the interaction between these structured slow sets and input-dependent reconfiguration of the high-dimensional vector field. Inputs transiently redirect trajectories through state space, whereas the slow sets constrain and temporally organize their subsequent evolution, enabling interval encoding, memory and reproduction. The extent of the slow sets determines generalization beyond the training range, and minimal dynamical systems endowed with analogous geometry reproduce the essential features of interval timing. Together, our findings identify structured slow transients as the dynamical substrate of temporal computation and provide a mechanistic interpretation of slow low-dimensional manifolds as emergent consequences of near-bifurcation state-space organization.

## Results

### Slow transient dynamics guide task-relevant trajectories despite attractor diversity

We trained recurrent neural networks to perform a flexible interval-reproduction task in which the temporal separation between two input stimuli (*S*_1_ and *S*_2_) defined a target interval *T* (Fig. 1a). After a variable delay period (*T*_*delay*_), an external *Go* cue instructed the network to reproduce the stored interval. The network output, computed as a weighted sum of unit activities, was trained to generate a binary response precisely when the reproduced interval had elapsed. We considered both gated recurrent unit (GRU) networks (Cho et al., 2014; Jordan et al., 2021), which are a simplification of long short-term memory (LSTM) networks (Hochreiter and Schmidhuber, 1997), and classical vanilla recurrent neural networks (RNNs) (Elman, 1990; Sussillo and Barak, 2013) spanning a range of network sizes (see Methods). Across architectures, initializations and network sizes, approximately 110 independently trained networks successfully learned the task, with larger networks exhibiting lower training errors and reduced variability across independent trainings (Supp. Fig. S1a, b). Consistent with previous studies, population activity evolved along low-dimensional trajectories that were largely conserved across network sizes and architectures, suggesting that successful timing relies on common dynamical principles despite substantial differences in network implementation (Figure 1b; Supp. Fig. S1c). Despite exhibiting nearly identical performance and similar low-dimensional trajectories, independently trained networks converged to fundamentally different state-space representations, as seen by the number of fixed points identified by convergence of randomly initialized trajectories in state space (Fig. 1b).

**Figure 1:**
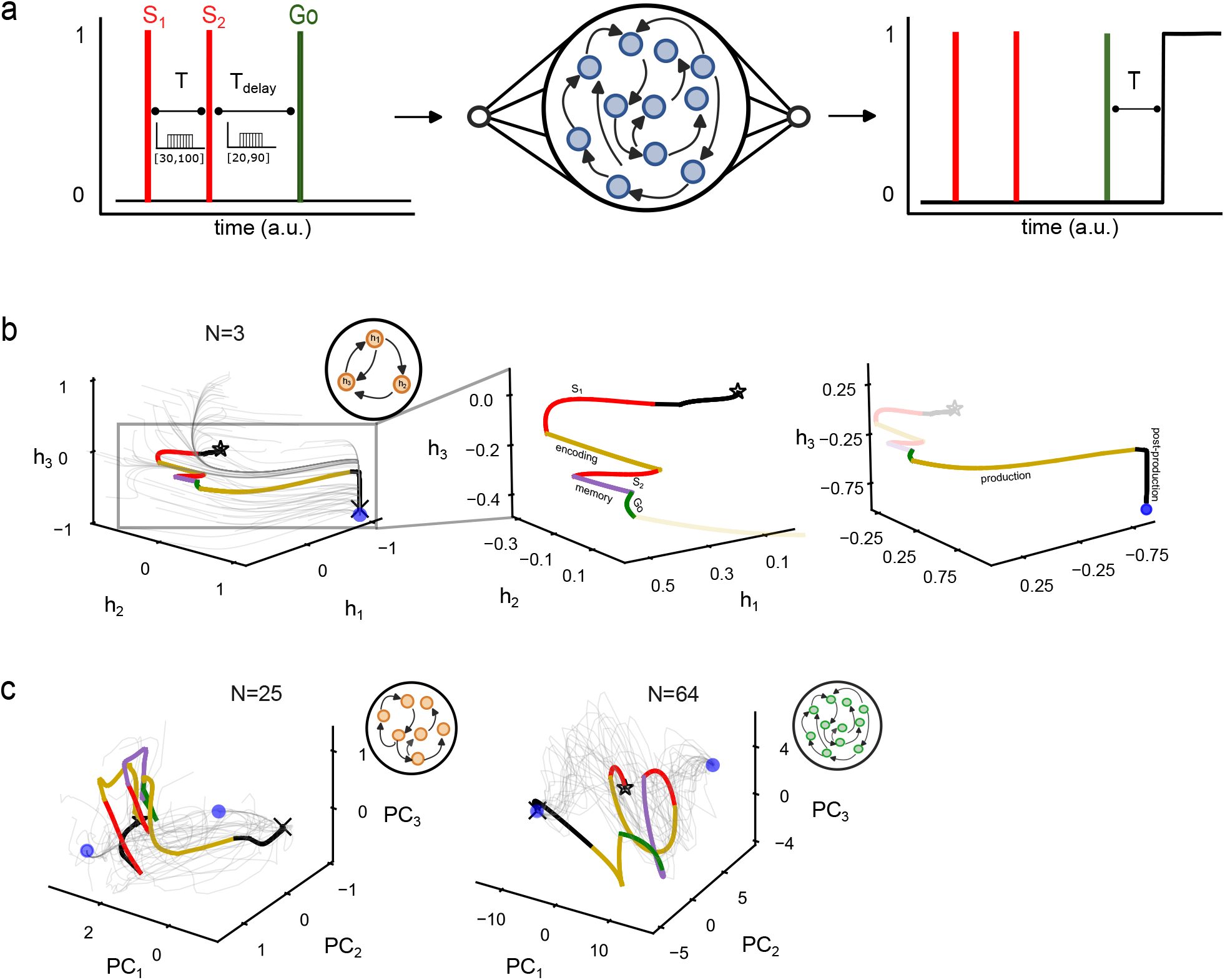
Equivalent low-dimensional dynamics of diverse trained RNNs. (a) Schematic representation of the interval production task. The interval *T* determined by *S*_1_ and *S*_2_ is kept during a variable *T*_*delay*_, and reproduced upon a *Go* cue. Exemplary task-relevant population activity trajectories in state-space for a (b) *N* = 3 gated recurrent unit (GRU) network (including zoomed insets) and (c) PCA space for *N* = 25 unit GRU or *N* = 64 node vanilla RNN. In the trajectories, red lines denote *S*_1_ and *S*_2_ presentation, green - the *Go* cue, yellow - encoding and reproduction of *T*, purple - trajectory during *T*_*delay*_. Star/cross: start/end of the trajectory; circle: stable fixed point. Insets in (b,c): schematics of the respective RNN.

To characterize this diversity formally, we analyzed ten independently trained 3-unit GRU networks (Supp. Fig. S1b) as tractable representative examples. We systematically identified their bifurcation structure, tracking how the number and stability of steady states changed as a function of the input amplitude. The diversity of fixed points solutions was also prevalent in the minimal networks - the autonomous dynamics (*input* = 0) spanned mono-, bi-, or multistable regimes, whereas other networks operated in the immediate vicinity of saddle-node (*SN* ) bifurcations (Fig. 2a; Supp. Fig. S2a). Thus, successful interval reproduction was compatible with a wide range of attractor architectures, demonstrating that learning does not converge onto a unique attractor solution. Task-relevant trajectories however revealed a second, more striking feature. During interval encoding, memory and reproduction, trajectories did not reside near, are often did not converge toward the attractors identified by the bifurcation analysis (compare yellow and purple parts of the trajectory in Fig. 1b with Fig. 2b, Supp. Fig. S2b-d). Instead, activity was continuously maintained far from the asymptotic fixed points of the system during the task execution. This observation suggested that timing is implemented primarily through transient dynamics rather than attractor convergence.

**Figure 2:**
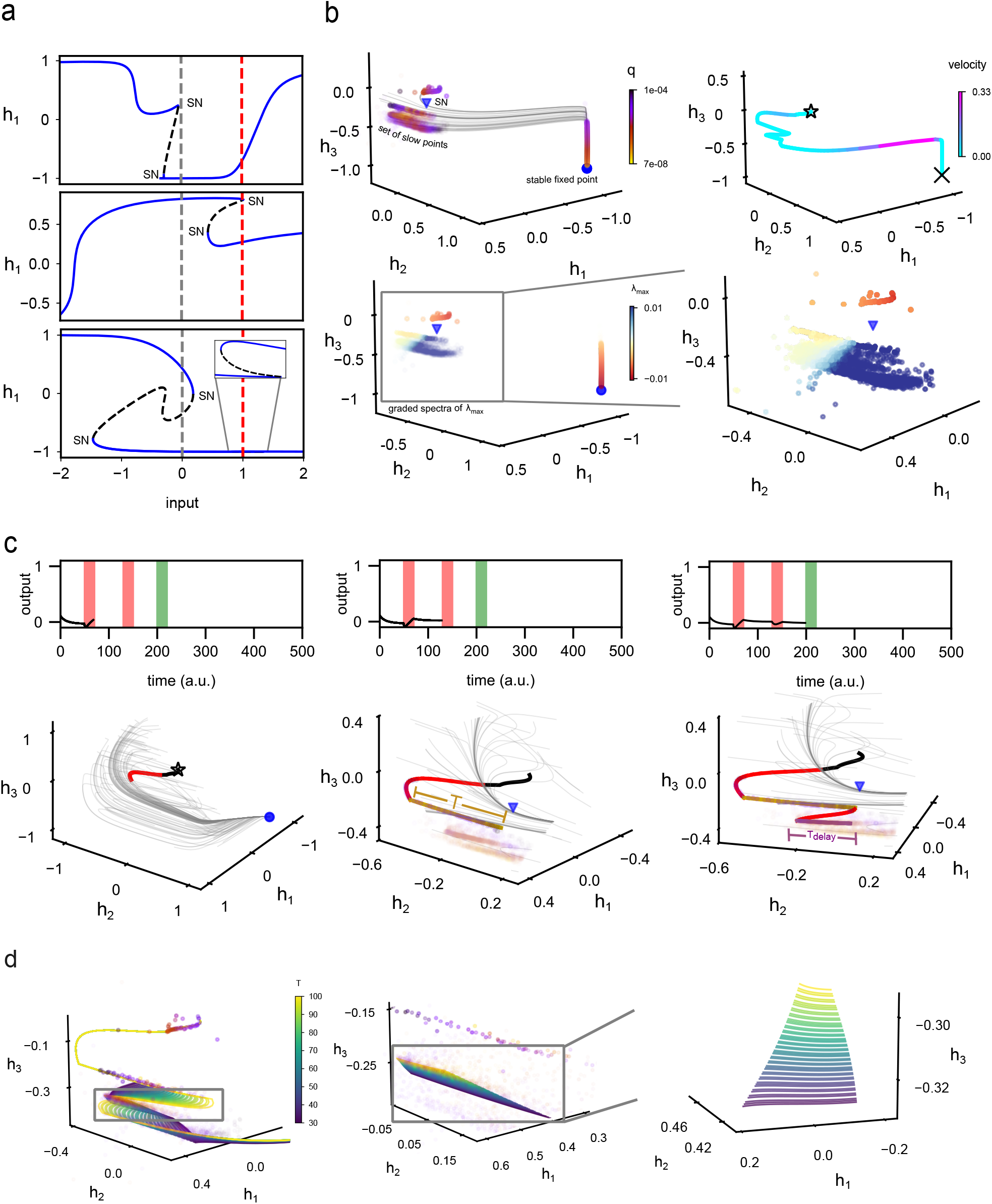
Structured transient dynamics implement interval timing in trained RNNs. (a) Bifurcation diagrams of three exemplary trained *N* = 3 GRU networks. Blue / blacks lines: stable / unstable fixed points. SN: saddle-node bifurcation. Gray/red dashed lines: GRU solutions for *input* = 0 / *input* = 1. (b) Characterization of the state space during computations for the network depicted in (a, top) / Fig.1b, top row. Top row: The set of slow points/fixed points identified using the norm of the dynamics, *q*, for *input* = 0, including the flow visualized through randomly initialized trajectories (grey) is shown on the left, and the task relevant trajectory color-coded by its velocity in state space shown on the right. Circle / triangle: position of the fixed point / saddle-node bifurcation corresponding to (a). Bottom row: Maximal local eigenvalue (*λ*; zoom inset to the right) of the corresponding state-space structures. Remaining symbols as in Fig.1. For other examples see also Supp. Fig. S2. (c) Top: Snapshots of the temporal evolution of the network’s output for the example in (b) during *S*_1_ (left), in the *T* -encoding period (middle) and during *T*_*delay*_ (right). Bottom: corresponding task-relevant trajectory segments and flow visualization. Remaining symbols and color coding as in (b). (d) Task-relevant trajectories for different *T* s for the example in (b), including zoomed representations (insets) of the trajectories’ segments during the delay period.

To identify the structures governing these transients, we analyzed the auxiliary function 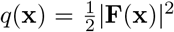. The minima of this function correspond to the fixed points of this system, and regions with *q ≈* 0 reveal sets of slow points and quasi-stable dynamics (Sussillo and Barak, 2013; Koch et al., 2024; Zheng et al., 2025; Koch and Nandan, 2026). Despite their diverse attractor architectures, all trained networks shared a common dynamical feature: extended *organized sets* of slow points (the criticality case in shown in Fig. 2b; other trained network examples in Supp. Fig. S2c, d). These slow sets exhibited graded spectra of maximal local eigenvalues (*λ*_*max*_) spanning from weakly negative to weakly positive. This organization of the eigenvalues generated two complementary effects. First, it dramatically slowed the flow of activity, preventing rapid collapse onto stable fixed points during encoding and memory (Fig. 2b - top right; Supp. Fig. S2e). Second, it organized state-space flow along a dominant direction. Sample trajectories initialized throughout state-space converged toward these regions and subsequently evolved along them (Fig. 2b -top left; Supp. Fig. S2c,d -left), indicating that slow sets act as *dynamical scaffolds* that constrain transient evolution. Thus, although attractor architectures differed markedly across networks, the organization of slow transient dynamics was highly conserved.

We next explain how the temporal evolution of the trajectory is regulated on the representative network poised near a saddle-node bifurcation (Fig. 2a, top). The first stimulus, *S*_1_, transiently stabilized an input-dependent fixed point towards which the flow was transiently redirected. Although the stimulus was too brief for convergence to the attractor, following the stimulus removal, the activity trajectory was displaced along the organized slow set. Hence, between the stimuli the activity evolved along this slow scaffold rather than collapsing toward the baseline attractor, such that the encoded interval *T* corresponded to the distance traversed through the slow region(Fig. 2c, left and middle). The second stimulus, *S*_2_, again transiently reconfigured the vector field and its removal resulted in repositioning the activity to a different location on the same slow set (Fig. 2c, right; Supp. Fig. S2b). Because the slow set exhibits a graded spectrum of near-zero eigenvalues, the parts of the trajectory initiated from different locations evolved with appropriately matched temporal profiles. Memory was therefore maintained as a transversal along the structured slow set, similarly to the *T* encoding, rather than as persistence at a stationary state. Following the *Go* cue, activity continued to evolve along the same scaffold and eventually leaving it, reproducing the stored interval without requiring convergence to an attractor (Supp. Fig. S2b).

To obtain a global view of the computation, we reconstructed the extended low-dimensional structures traversed by the trajectories across task conditions (Supp. Fig. S3a). Consistent with previous studies, the population activity remained confined to low-dimensional manifolds. However, our analysis indicates that these manifolds only describe where the activity lies, but do not mechanistically explain the computation. Computation emerged from the interaction between input-dependent reconfiguration of the vector field, and the structured slow sets that regulate the transient evolution of population trajectories (Supp. Fig. S3b,c), thereby transforming inputs into temporally controlled computations. Hence, different intervals corresponded to distinct traversals on a common slow scaffold, whose graded spectrum of local eigenvalues determined the pace of temporal evolution required for interval encoding, memory and reproduction (Fig. 2d). Importantly, the longest intervals were associated with the smallest eigenvalue magnitudes, corresponding to the slowest decay modes of the scaffold, thereby linking temporal scaling directly to the spectral structure of the underlying dynamics (compare Fig. 2d with Supp. Fig. S3a). These findings therefore identify a common dynamical mechanism underlying otherwise diverse trained solutions: organized sets of slow points with characteristic eigenvalue ordering regulate the temporal pace of population trajectories and provide a candidate state-space substrate for temporal computation.

### Learning drives networks toward near-critical regimes generating slow sets

We next asked how the conserved slow set structures identified above emerge during learning. Tracking the evolution of network dynamics throughout training, as exemplified by the GRU network organized at criticality (Fig. 2a, top), revealed a progressive reorganization of the underlying state-space. Early in training, trajectories rapidly collapsed onto the stable attractor. This single state-space structure hinders responsiveness to the task-related signals and reduces separability between different task phases, both of which are necessary for temporally controlled progression. As a result, timing performance was poor (Fig. 3a and b, left column). As learning proceeded, the local eigenvalue spectrum progressively compressed toward zero, giving rise to extended sets of slow points. The trajectories became increasingly confined to coherent flow lines, producing a marked increase in flow directionality (i.e. between epochs 6 and 21, Fig. 3b) and a corresponding improvement in task performance. In networks for which complete bifurcation analyses could be performed, this reorganization coincided with the emergence of *SN* bifurcations and the positioning of the network’s parameters in their immediate vicinity (Fig. 3a; additional example shown in Supp. Fig. S4a,b). Thus, learning progressively transformed attractor-dominated dynamics into organized transient flow in state space.

**Figure 3:**
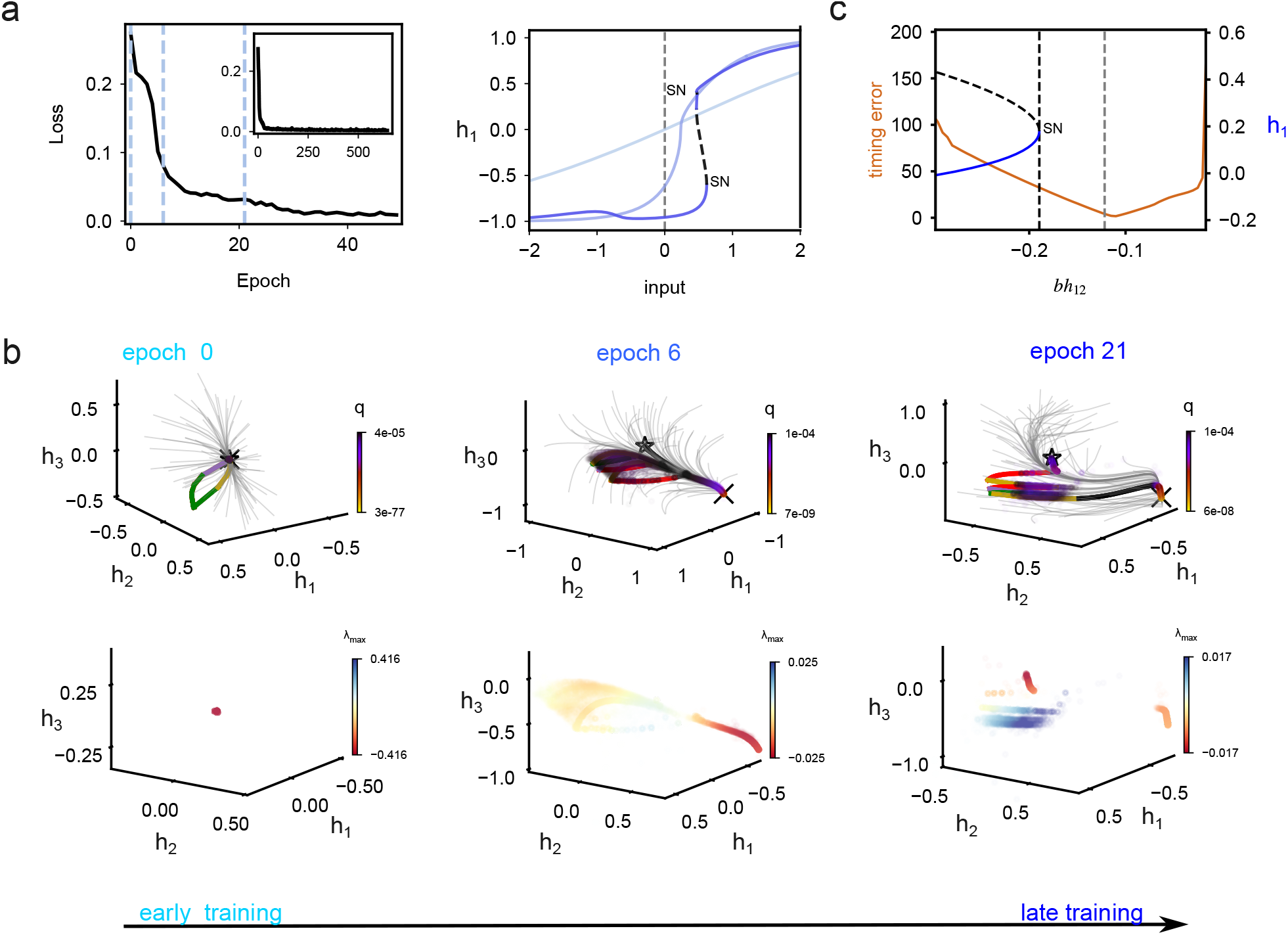
Emergence of sets of slow points during GRU training. (a) Representative loss curve of the initial 50 trained epochs with a zoomed inset of the full curve (left) and the evolution of the bifurcation structure of the network during training for the marked epochs (0, 6 and 21, right). Blue / black lines: stable / unstable fixed points; gray dashed line: organization at *input* = 0. The example corresponds to Fig. 2a, top. (b) Snapshots of the task-relevant trajectories and the sets of slow points (top), as well as the corresponding local maximal eigenvalue (bottom) at the marked epochs during training. Trajectory color-coding as in Fig. 1b. For additional example, see Supp. Fig. S4a,b. (c) Bifurcation diagram of a trained 3-unit GRU network with respect to a bias parameter *bh*_12_ (line color as in (a)) and the respective timing error calculated for different *bh*_12_ values (red line). Vertical black/grey dashed line: position of the *SN* bifurcation / organization after training. Related to Supp. Fig. S4c.

This observation closely matches a canonical phenomenon from dynamical systems theory. Systems positioned near saddle-node bifurcations generate extended regions of slow dynamics, termed ghost sets, with graded spectra of near-zero eigenvalues (Strogatz and Westervelt, 1989; Koch et al., 2024; Koch and Nandan, 2026). The learned slow set structures exhibit the same signatures, including flow slowing and directional funneling (Fig. 2b, 3b, bottom row). Hence, learning did not encode temporal intervals through stable states, but by creating state-space geometries that support structured transient evolution of the population activity. Although in many networks the bifurcations were induced by the external inputs, in some of the trained examples similar near-critical organization emerged with respect to the internal network parameters (i.e. the bias parameter *bh*_12_, example shown in Fig. 3c; Supp. Fig. S4c -middle column). This observation enabled us to directly test the functional importance of near-critical organization by perturbing a single bifurcation parameter while keeping all remaining parameters fixed. Timing performance was maximal in the immediate vicinity of the bifurcation and deteriorated as the system moved away from the critical regime (Fig. 3). This loss of performance coincided with the reorganization or disappearance of the ghost set (Supp. Fig. S4c). In training examples where the parameter responsible for critical organization could not be identified due to the high dimensionality of the system, we nevertheless identified slow sets characterized by an equivalent eigenvalue structure (examples in Supp. Fig.S2c,d). Although we cannot exclude the possibility that different mechanisms may give rise to these structures, the existing literature has so far associated them primarily with ghost sets Koch et al. (2024); Koch and Nandan (2026). These observations therefore indicate that organized slow transients are functionally necessary for temporal computation rather than epiphenomenal consequences of training. Furthermore, they also imply that positioning in a critical vicinity of the bifurcation where the ghost set exists, after the coalescence of the stable and saddle nodes, is crucial for controlling the temporal evolution of the trajectories during task evolution.

### The extent of the slow sets determines generalization to untrained intervals

We next asked whether the extent of the slow sets determines the networks’ ability to generalize. Here, we define generalization as the ability of a trained network to accurately perform task conditions that were never encountered during training. Because the ghost slow sets not only generate long timescales but also funnel trajectories along task-relevant directions, we hypothesized that their extent constrains the range of computations that can be realized. To test this idea, we trained a 3-unit GRU network to memorize and reproduce a single interval (*T* = 80a.u.; corresponding *T*_*delay*_ = 50a.u.). As expected, when evaluated using a different interval (*T* = 60a.u.), the network failed to reproduce the target duration accurately (Δ*T* = *−*18a.u.; Figure 4a), indicating that successful extrapolation does not arise trivially from training. Nevertheless, the trained network exhibited organization near a *SN* bifurcation and an associated ghost slow set (Fig. 4b; Supp. Fig. S6a,b). Our previous analysis suggests that different intervals correspond to different positions along a common slow set, with local geometry determining the effective pace of subsequent evolution (Fig.2d). We therefore asked whether repositioning the trajectory along the existing slow set could generate intervals that had never been explicitly learned. Remarkably, varying the amplitude of the *Go* cue enabled the same network to reproduce a continuous range of intervals (*T ∈* [20, 100]) despite having been trained on a single target duration (Fig. 4c).

**Figure 4:**
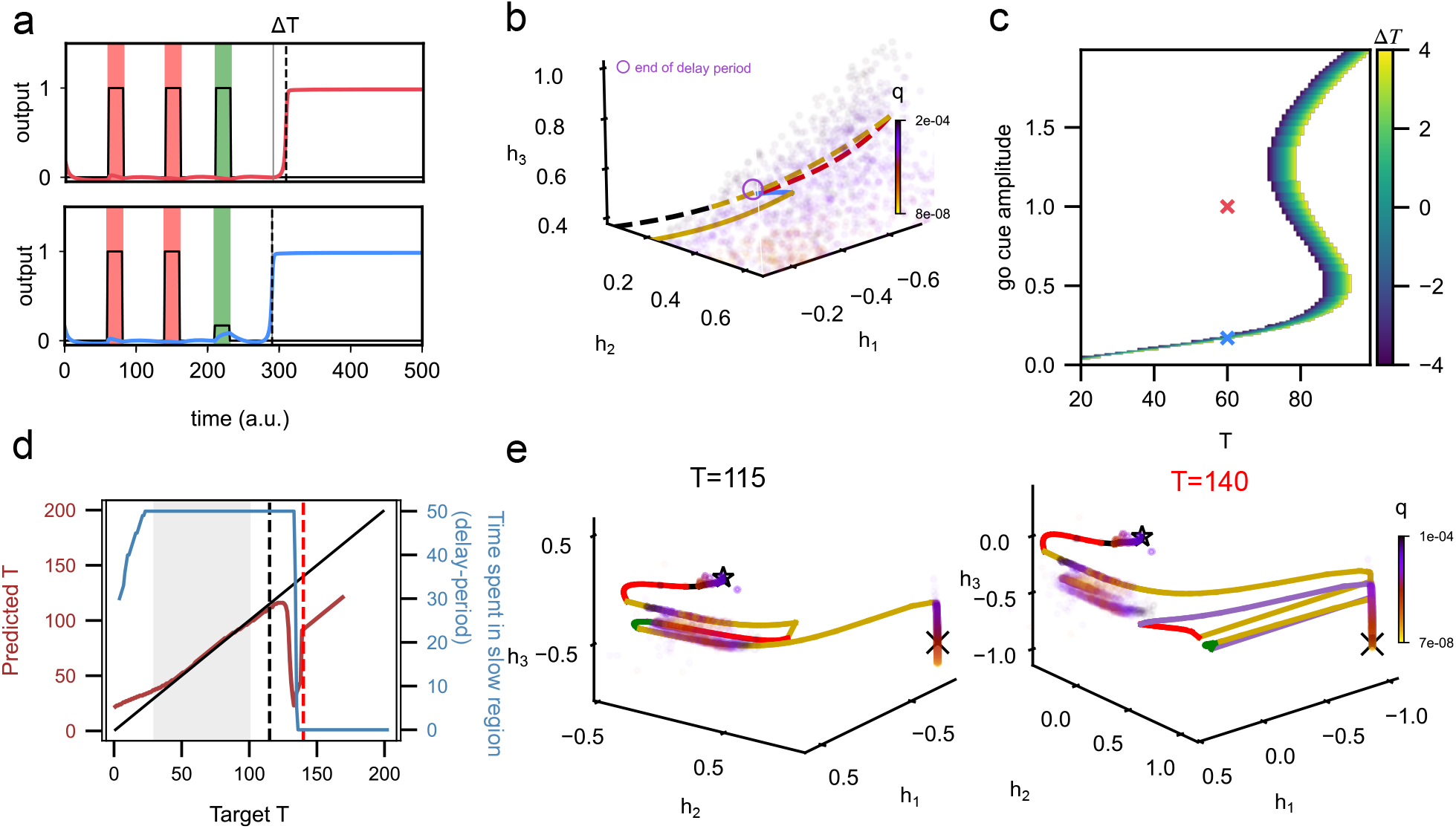
Generalization property of trained networks relies on the extent of the set of slow points. (a) Response timeseries of a *N* = 3 unit GRU network trained to encode *T* = 80a.u., and tested on *T* = 60a.u. when the *Go* cue amplitude is equal to 1 (top), and 0.172 (bottom - corresponding to blue cross in (c)). Solid black/cyan (red) lines: input/output timeseries, Δ*T* - timing error in response. (b) The set of slow points of the trained network characterized by *q* and the population activity trajectories for the two *Go* cue amplitudes shown in (a). Trajectories color coded as in Figure 1b. Dashed/solid lines correspond to *Go* cue amplitude 1*/*0.172 respectively. Related to Supp. Fig. S6a,b. (c) Identified *Go* cue amplitudes and corresponding Δ*T* quantification resulting in generalization for encoding 20 *< T <* 100. (d) Generalization extent of the network in Fig.2a -top. Solid black/red line: optimal / actual predicted *T* . Blue line: Time spent by the trajectory during the memory phase (delay period). Gray shaded area: *T* interval presented during network training. Black/red dashed lines correspond to *T* = 115*/*140a.u. respectively, shown in (e). Additional example shown in Supp. Fig. S6c,d. (e) Task-dependent trajectories and set of slow points for *T* = 115a.u. (left) and *T* = 140a.u. (right).

Moreover, if generalization is an emergent consequence of the slow set extent, then networks endowed with larger and more smoothly organized ghost sets should generalize more broadly. To test this prediction, we analyzed networks trained on intervals uniformly sampled from *T ∈* [30, 100) and *T*_*delay*_ *∈* [20, 90) (examples from Fig.2a,top). Although independently trained networks exhibited substantial variability in out-of-training performance, their generalization capabilities were strongly constrained by the extent of their slow sets (Fig. 4d,e; Supp. Fig. S6c,d). Networks generalized precisely over those intervals for which trajectories remained confined to the slow set during encoding and memory, whereas performance deteriorated once trajectories exited this region during these task phases. These results identify a direct relationship between state-space geometry and computational generalization. Generalization is therefore not an additional feature acquired through exposure to multiple intervals, but an intrinsic consequence of the extent of the slow sets created during learning. This determines the range of temporal computations that a network can perform beyond its training experience.

### Ghost sets are sufficient for flexible temporal computation

To test whether the structured slow sets uncovered in trained networks constitute a minimal dynamical ingredient for flexible temporal computation, we constructed a low-dimensional model with three variables (*G*_1_, *G*_2_, *G*_3_) whose dynamics is characterized by individual ghosts used for coordinating the different responses, a memory variable (*z*), and a timing variable (*x*; Figure 5a, details in Methods), inspired by the models of neural integrators that predict time Durstewitz (2003). The first input pulse activates *G*_1_ and initiates a gradual increase of *z*. When the system receives the second pulse while *G*_1_ is still active (through its ghost), *G*_2_ will be activated, terminating the accumulation of *z* and thereby storing the encoded interval. The system then remains in a waiting state until a third pulse activates *G*_3_, which resets the ghost variables and initiates the evolution of *x* through another ghost set, in this case ghost of a line attractor. Critically, the extent of this slow region scales linearly with *z*, such that longer intervals induce longer traversals through the ghost set. The resulting transient duration of *x* directly reproduces the stored interval without learning or parameter optimization (an exemplary response is shown in Figure 5b). The model thereby autonomously generates the encoding, memory, and correct timing dynamics, accurately reproducing target intervals (error *<* 10%) across a broad range of *T*, despite never being explicitly trained for interval timing.

**Figure 5:**
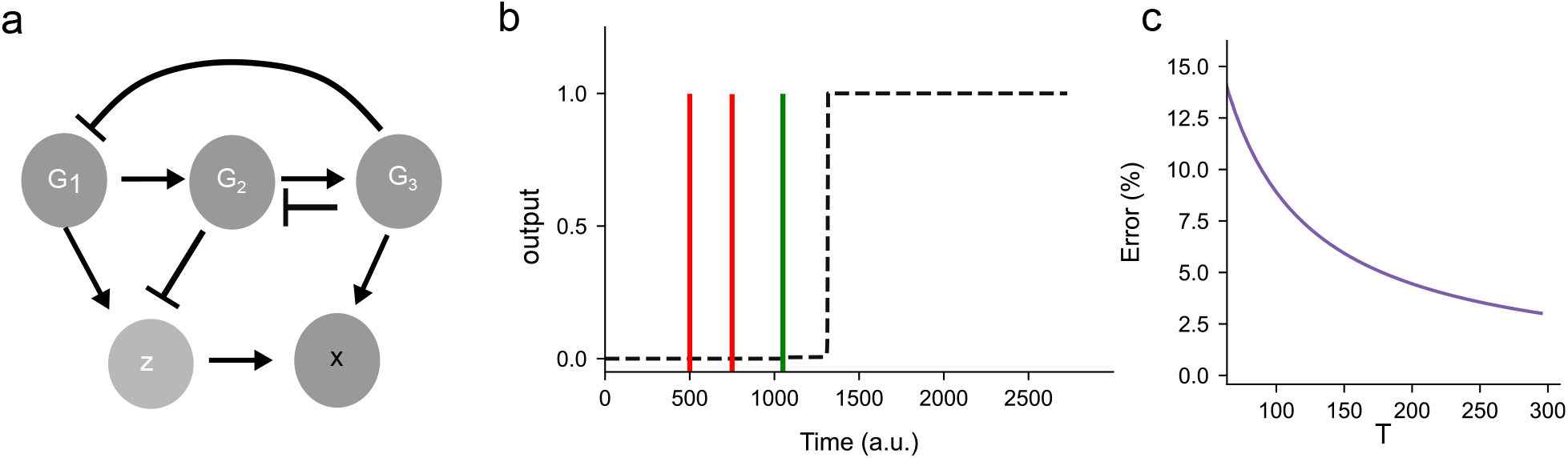
Dynamical systems model for flexible temporal computations. (a) Schematic representation of the model: *G*_1_ *− G*_3_ - single ghost variables, *x* - timing variable, *z* - memory variable (details in Methods). Arrows/block arrows: activation/inhibition. (b) Exemplary time series depicting the model response for *T* = 250a.u. Black dashed line: *dx/dt* as a function of time. (c) Estimated *T* interval range for which the model achieves *<* 15% error in prediction.

Thus, ghost-like slow sets are not merely an emergent property of trained recurrent networks, but constitute sufficient dynamical ingredients for temporal computation. More generally, temporal computation arises from the structured organization of transient flows in state space, linking near-critical dynamics, neural manifolds, and computation. We therefore argue that such minimal models provide useful theoretical benchmarks for temporal computation by isolating the essential dynamical ingredients required for timing and enabling direct comparison with biological population activity and trained recurrent neural networks.

## Discussion

Flexible behavior requires neural activity to evolve through precisely organized transient dynamics. Although low-dimensional neural manifolds have been widely observed across cognitive tasks Remington et al. (2018); Wang et al. (2018); Egger et al. (2019); Langdon et al. (2023); Perich and Gallego (2025), the dynamical mechanisms that generate and constrain the evolution of the trajectories along these manifolds have remained unclear. Here we show that recurrent neural networks trained on interval timing converge to diverse attractor configurations, however, they exhibit a conserved organization at the level of transient dynamics. Across independently trained solutions, the temporal evolution of the trajectories is consistently organized by structured ghost sets characterized by graded spectra of near-zero eigenvalues that constrain and channel the population activity flow. These structures emerge during learning through positioning the networks in a critical vicinity of saddle-node bifurcations. Recent work has shown that near-critical dynamics in neuronal and artificial networks exhibit long timescales Levina et al. (2007); Deco and Jirsa (2012); Shew and Plenz (2013); Ma et al. (2019); Kelso (2023); Goto et al. (2025); Rossi et al. (2025); Hancock et al. (2025), high-dimensional global modes, and abrupt learning transitions Pachitariu et al. (2026); Dinc et al. (2026), suggesting a link between criticality and computational capability. We have extended these findings by identifying structured slow sets as explicit geometric objects that not only organize temporal computations, but their extent also predicts the generalization to unseen intervals. Our results also provide a mechanistic reinterpretation of low-dimensional neural manifolds. Although neural activity remains confined to low-dimensional trajectories during task execution Wang et al. (2018); Remington et al. (2018); Egger et al. (2019); Jazayeri and Ostojic (2021), we demonstrate here that these manifolds do not themselves determine computation. Instead, computation emerges from the interaction between input-driven reconfiguration of the high-dimensional vector field and the existing slow set that governs the subsequent evolution of the population dynamics. Different temporal intervals correspond to distinct traversals of a shared slow scaffold, whose eigenvalue structure determines the rate of progression through state space. Thus, low-dimensional slow manifolds reflect constraints imposed by dynamical organization rather than serving as primary computational substrates.

A central implication of this view is that task-relevant information is encoded and transformed through evolution along the structured slow regions, rather than by position on a stable attractor. We further demonstrate the sufficiency of this mechanism using a minimal dynamical system endowed with analogous slow set geometry. Without learning or parameter optimization, the system reproduces interval timing across broad intervals range via traversal of ghost-like slow regions whose spatial extent governs the duration of transient evolution. This establishes structured slow sets as a minimal dynamical ingredient for temporal computation. Importantly, our findings suggest a constructive principle for designing efficient and minimal artificial neural networks: embedding extended slow set regions Schmidt et al. (2021); Durstewitz et al. (2025); Dinc et al. (2026) should improve robustness and generalization in temporal tasks. Finally, this framework yields experimentally testable predictions. During learning, neural population activity should progressively align with slow modes corresponding to task variables. Perturbations would induce rapid relaxation along fast directions while preserving constrained evolution within the slow scaffold. Moreover, the spatial extent of the slow regions should predict behavioral generalization beyond training conditions. These predictions are directly testable from neural recordings.

Taken together, the presented results unify diverse observations of low-dimensional neural dynamics under a single mechanistic account grounded in state-space organization, showing that temporal computation emerges from underlying dynamical structure rather than specialized or static representations, and identifying near-critical slow dynamics as a principle for flexible temporal computation.

## Data availability

Codes for generating numerical results are available in https://github.com/gayathriramesan/slow_points_for_timing

## Author contributions

Conceptualization, A.K; methodology, A.K., G.R., A.N., D.K.; investigation, G.R, A.N., D.K.; writing – original draft, A.K. and G.R.; writing – review and editing, all authors; visualization, G.R.; supervision, A.K.

## Acknowledgments

A.K. acknowledges funding by the Lise Meitner Excellence Programme of the Max Planck Society, D.K—EMBO Fellowship (Grant No. ALTF 310-2021), A.K. is a member of the Nordrhein-Westfalen (NRW) network iBehave. We thank U. Feudel, P. Ashwin and the CCL group members for valuable discussions.

## Materials and methods

### Network structure and training

We considered two different classes of recurrent neural networks: a gated recurrent neural network Jordan et al. (2021) and a vanilla RNN Elman (1990); Sussillo and Barak (2013). The gated recurrent unit dynamics is given by:

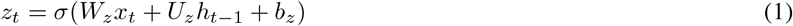

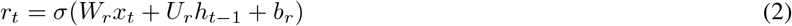

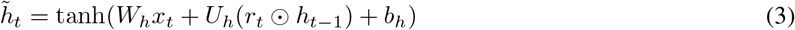

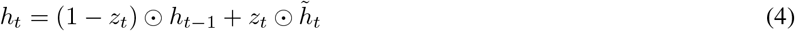

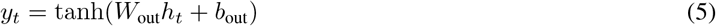

where *x*_*t*_ is the input, *h*_*t*_ - hidden state, *z*_*t*_ - update gate, *r*_*t*_ - the reset gate, *σ*(*·*) denotes a sigmoid activation function, *⊙* denotes element-wise multiplication, and tanh(*·*) is applied element-wise. For most of the analysis in the manuscript we refer to the continuous-time version of the GRU obtained by Euler discretizations of 4. For further details on the complete discretizations, refer to Jordan et al. (2021).

The dynamics of the Vanilla RNN model is given by:

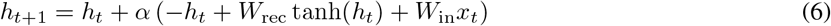

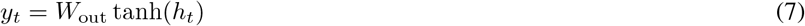

where *α* = *dt/τ, x*_*t*_ is the input, *h*_*t*_ - the hidden state, *W*_in_ are the input weights, *W*_rec_ are the recurrent weights, and *W*_out_ are the output weights.

We trained 10 GRUs each of 3,5,10..50 nodes and 10 Vanilla RNNs of 64 nodes. All networks were trained using classical backpropagation through time in TensorFlow. https://github.com/gayathriramesan/slow_points_for_timing. The loss function for training was computed as the mean squared error between the target and the prediction of the trained network:

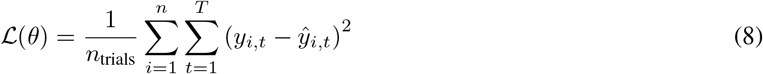

where *y*_*i,t*_ refers to the *i*^th^ prediction of the trained network at time *t*, and *ŷ*_*i,t*_ the corresponding target. The network was trained using the Adam optimiser with a learning rate of *η* = 0.01, *β*_1_ = 0.9, *β*_2_ = 0.999. The networks were trained for a maximum of 5000 epochs. The training was additionally halted if the loss does not improve for 1000 consecutive epochs, and the best weights were then restored. The network receives the first input *S*_1_ always at *t* = 50a.u. The time interval between the first two pulses *S*_1_ and *S*_2_, *T*, and between *S*_2_ and the ‘Go’ cue, *T*_delay_, were randomly sampled from the following distributions: *T ∼ U* [30, 100), *T*_*delay*_ *∼ U* [20, 90).

### Identifying state-space structures

The attractor landscapes of the trained networks were identified for the continuous-time ordinary differential equations of the trained networks using the *XPPAUT* bifurcation analysis package (Ermentrout and Mahajan, 2003). The results were additionally verified using the *BifurcationKit* package in *Julia* (Veltz, 2020). For all the randomly initialized trajectories depicted as grey lines in the figures, the trained networks were numerically integrated using the Euler scheme in Python.

Slow points in state space were identified by minimizing the scalar function *q* Sussillo and Barak (2013) *using SciPy’s minimise* package with Newton’s method and a conjugate gradient solver. The trajectories of the RNN while performing the task were used as initial conditions for the identification of the slow points so that the solutions remain bounded within the state space. A *q*_threshold_ of 0.0001 was generally used for the identification of the slow point. In the regions where the norm of a dynamical system is close to zero *q*(*x*) *< q*_*thresh*_, local linear expansion of the dynamics is still valid. Thereby the local eigenvalues were calculated by numerically evaluating the Jacobian at these points. For more details refer to (Koch et al., 2024).

The slow-set scaffold depicted in Figure 2d, Supp. Fig. S3 was identified by initializing the minimization of *q* from random initial conditions throughout state space, in contrast to initializing it close to task-dependent trajectory. To characterize the geometry of the manifold of slow points, we fit a smooth surface to the population activity using a two-step procedure. First, principal component analysis was applied to the 3D point cloud to identify the two directions of greatest variance, projecting each point onto a 2D parametric coordinate system. This reduced the surface-fitting problem to a 3D *→* 2D mapping while preserving the dominant geometric structure of the set of slow points. Second, a Radial Basis Function interpolation with a thin-plate spline kernel was fit to map the coordinates back to the original 3D space, yielding the unique manifold containing the slow points in state space.

All of the numerical parameters and the corresponding equations are provided at https://github.com/gayathriramesan/slow_points_for_timing.

### Ghost timer

The model has five state variables: three single ghost units *G*_1_, *G*_2_, *G*_3_, a ghost of a line-attractor *x*, and an elapsed-time integrator *z*, governed by:

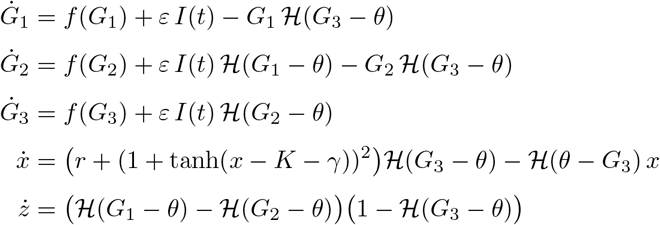

where *H* is the Heaviside function, *K* = *r*(*z* + 23.1), and *f* (*u*) = *a*(*u* + *x*_off_)^3^ + *b*(*u* + *x*_off_)^2^ + *c*(*u* + *x*_off_) + *d* is a ghost polynomial with parameters (*x*_off_, *a, b, c, d*) = (*−*2, *−*0.5, 2, 0, *−*4.7407). The remaining parameters are *r* = 0.02, *γ* = 1, *ε* = 5, and *θ* = 4.

---

**Supplementary Figure S1:**
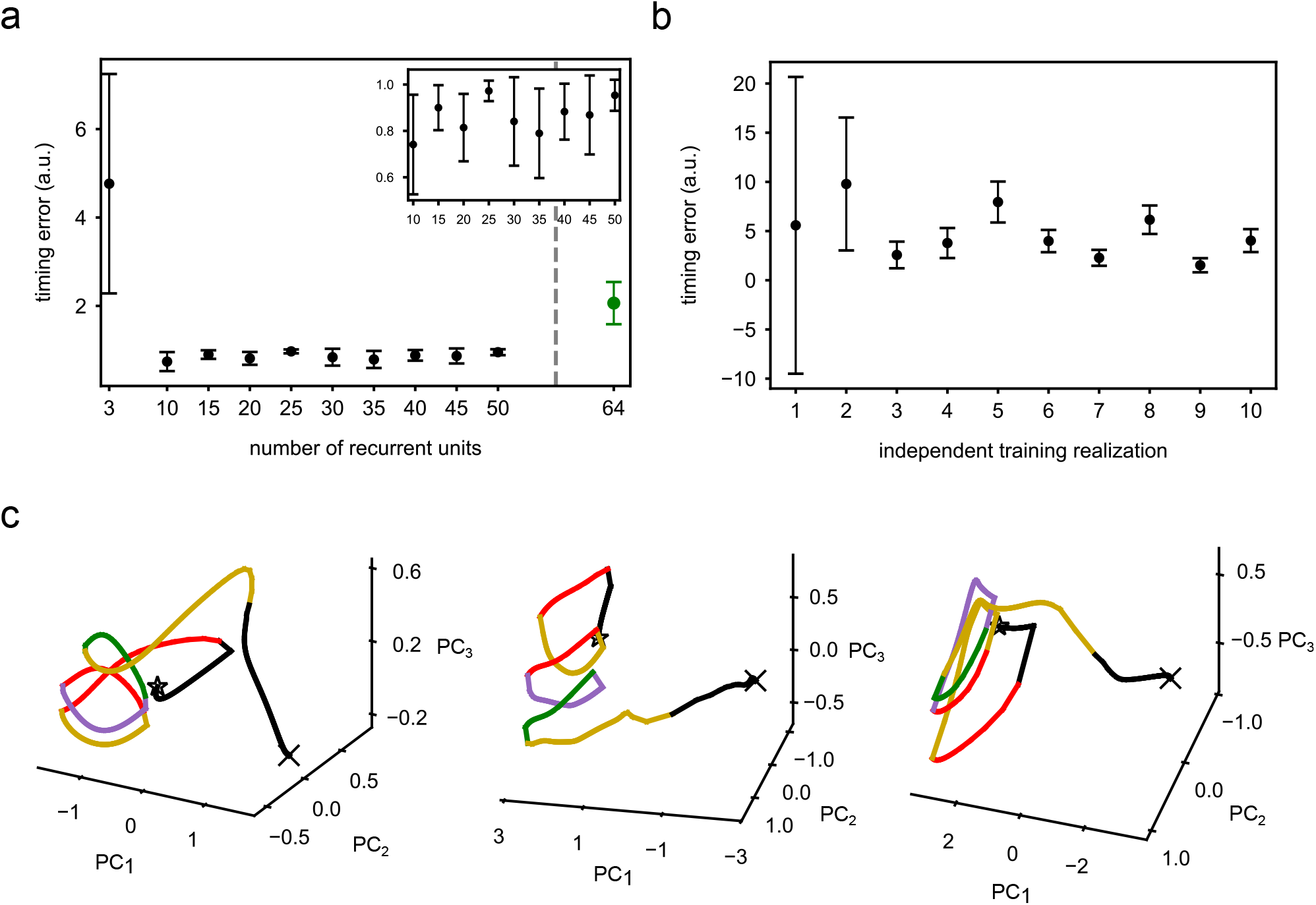
Dynamics of trained networks. (a) Timing error for trained GRU networks of 3, 10, 15, 20, 25, 30, 35, 40, 45 and 50*−* units, as well as 64*−* node vanilla RNN networks. Mean *±* standard deviation from 10 independent trainings in each case is shown. Inset depicts zoomed timing error for the GRU networks of size 10 *−* 50. (b) Timing error for the 10 independently trained 3*−* unit GRU networks is shown, with mean *±* standard deviation for 1000 different combinations of *T, T*_*delays*_. (c) Exemplary task-relevant population activity trajectories in PCA space for *N* = 10, 30, 50*−* unit GRU networks.

**Supplementary Figure S2:**
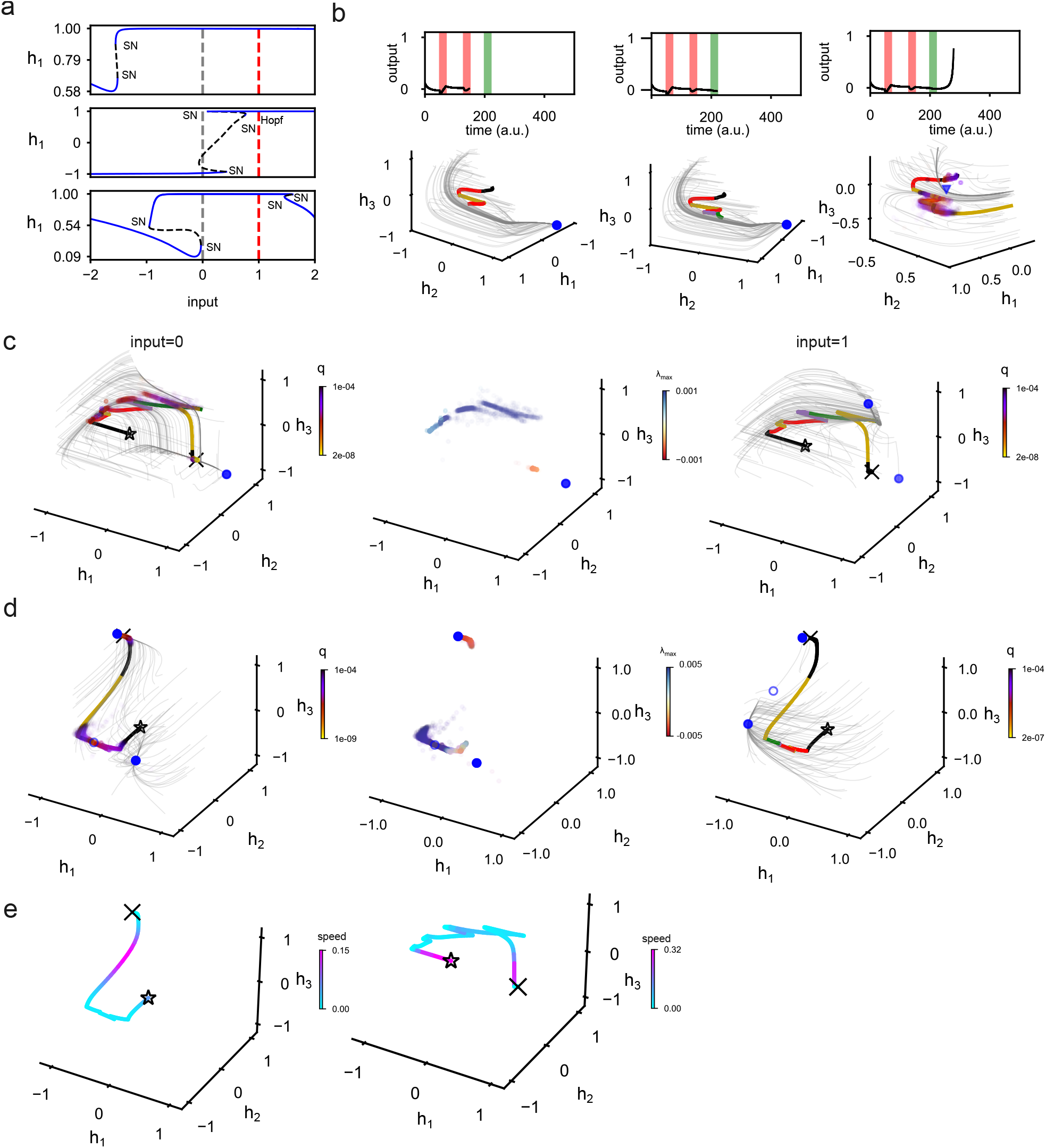
Characterizing transient dynamics of trained 3*−* unit GRU networks. (a) Bifurcation diagrams of three additional 3*−* unit GRU networks. *SN* - saddle-node bifurcation; *Hopf* - bifurcation; blue/black dashed lines: stable/unstable fixed points; grey/red dashed lines: organization for *input* = 0 / *input* = 1. (b) Remaining snapshots of the response timeseries and the temporal evolution of the state-space depicted in Fig. 2c. Color coding / symbols as in Fig. 2. (c) State-space characterization for the GRU networks in Fig.1a, middle. Left: Task-relevant trajectory and identified slow-point set / proxy of the flow (grey trajectories) for *input* = 0. Middle: Maximal local eigenvalues of the corresponding slow-point set. Right: State-space structure for *input* = 1, including the task-relevant trajectory. (d) Same as in (c) for Fig.1a, bottom. (e) Speed of the trajectories over time for (c) - left and (d) - right, respectively.

**Supplementary Figure S3:**
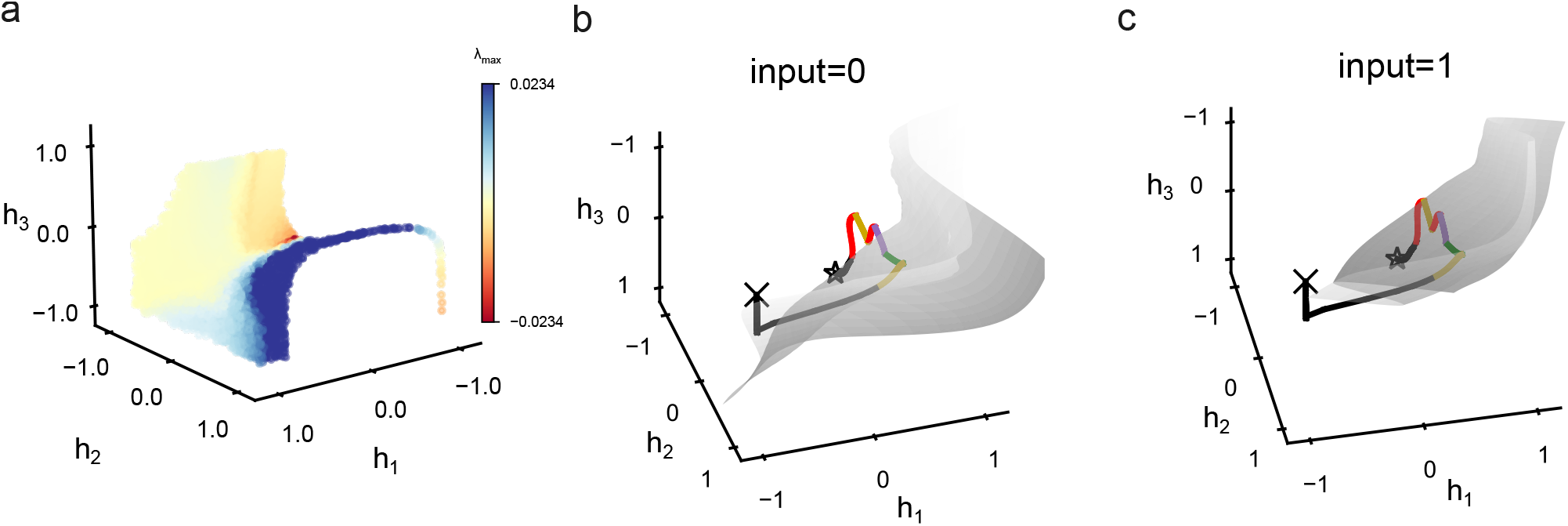
Computation between manifolds in state space. (a) Extended set of slow points corresponding to Fig.2c, color-coded by the maximal local eigenvalue. (b) 2D manifold representation and task-relevant trajectory at *input* = 0 and (c) *input* = 1. Trajectory color-coding as in Figure 1b.

**Supplementary Figure S4:**
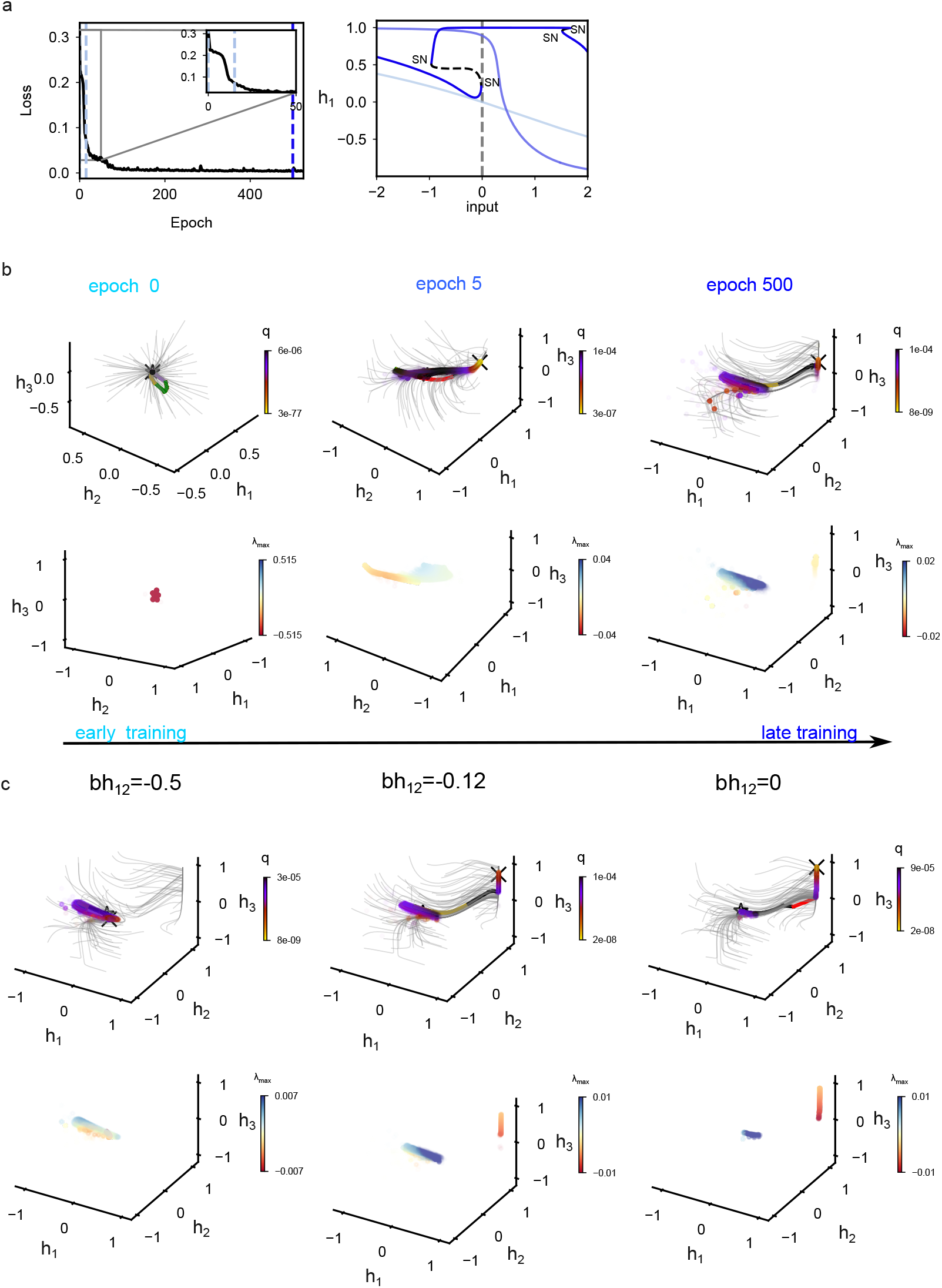
Sets of slow points are generated during network training. (a) Evolution of the bifurcation diagram (at epochs 0, 15, 500) and corresponding loss function during training for an additional 3*−* unit GRU network organized at criticality. (b) Corresponding state-space characterization (top) and evolution of the set of slow points depicted through the maximal local eigenvalue (bottom) for the network in (a). (c) Characterization of the state-space and the corresponding maximal eigenvalue of the detected sets of slow point for different parameter values of the network in Fig. 3c, for *bh*_12_ = *−*0.5 - left, *bh*_12_ = *−*0.12 - middle, and *bh*12 = 0 - right.

**Supplementary Figure S6:**
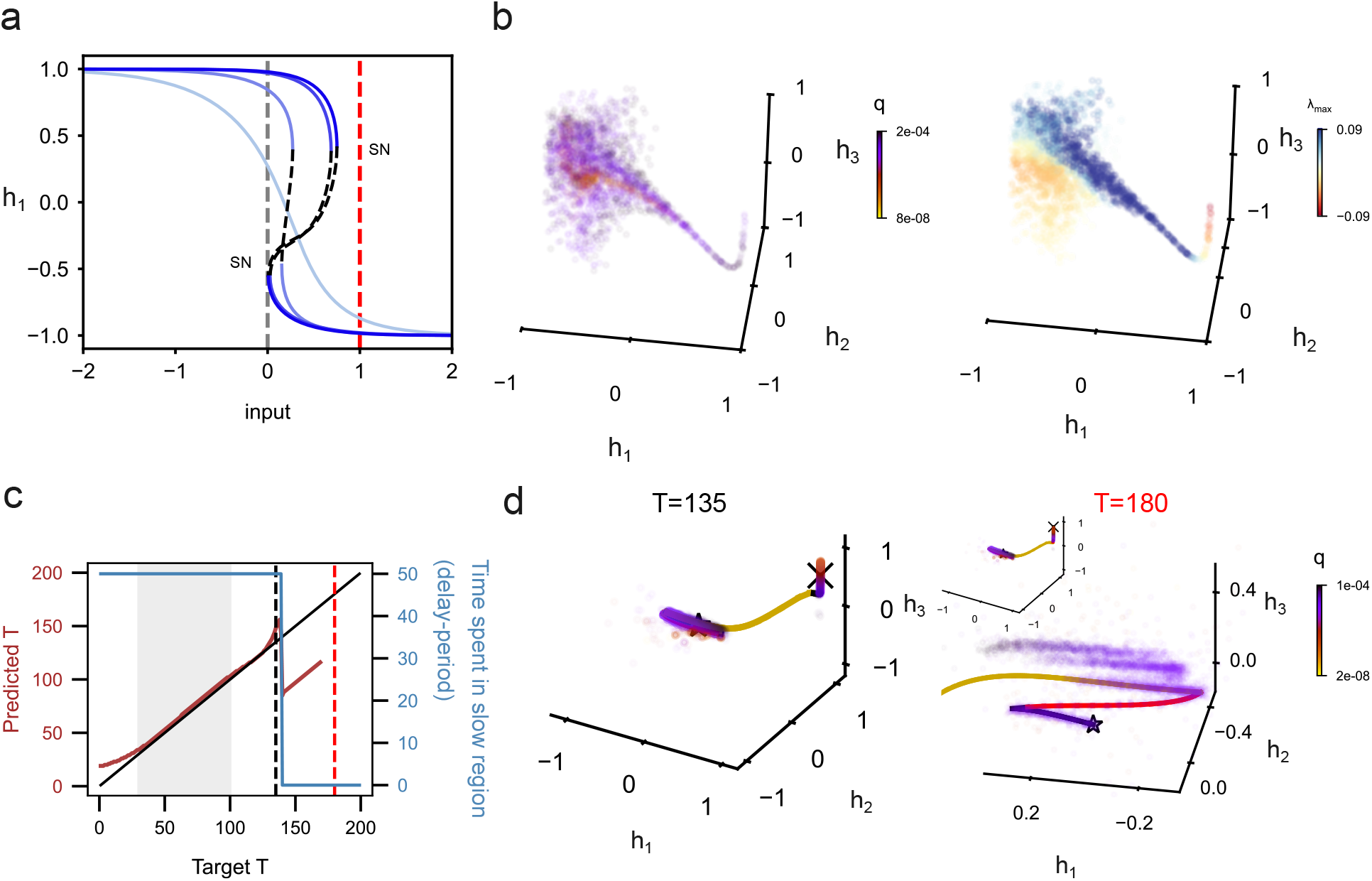
Sets of slow points underlie generalization to untrained *T* . (a) Evolution of the bifurcation diagram during training of the 3*−* unit GRU network depicted in Fig. 4a -top. Epochs 50, 100, 500, 1000, 4240 are shown. (b) Set of slow points and maximal eigenvalue characterization of the ghost set corresponding to (a) and Fig.4b. (c) Generalization properties of a trained 3*−* node GRU network shown in Fig.3c. Solid black/red line: optimal / actual predicted *T* ; dashed black/red line: tested *T* = 135*/*180a.u. depicted in (d). Blue line: Time spent by the trajectory during the memory phase (delay period). Gray shaded area: *T* interval presented during network training. (e) Task-dependent trajectories and set of slow points for *T* = 135a.u. (left) and *T* = 180a.u. (right).

